# Motor Cortex Excitability in Chronic Low Back Pain

**DOI:** 10.1101/2022.03.13.484179

**Authors:** E.J Corti, W Marinovic, A.T Nguyen, N Gasson, A.M Loftus

## Abstract

**Introduction:** Chronic pain is associated with dysfunctional cortical excitability. Research has identified altered intracortical motor cortex excitability in Chronic Lower Back Pain (CLBP). However, research identifying the specific intracortical changes underlying CLBP has been met with inconsistent findings. In the present case-control study, we examined intracortical excitability of the primary motor cortex using transcranial magnetic stimulation (TMS) in individuals with CLBP.

**Methods:** Twenty participants with CLBP (*M_age_* = 54.45 years, *SD_age_* = 15.89 years) and 18 age- and gender-matched, pain-free controls (*M* = 53.83, *SD* = 16.72) were included in this study. TMS was applied to the hand motor area of the right hemisphere and motor evoked potentials (MEPs) were recorded from the first dorsal interosseous muscle of the contralateral hand. Resting motor threshold (rMT) and MEP amplitude were measured using single-pulse stimulation. Short intracortical inhibition (SICI) and intracortical facilitation (ICF) were assessed using paired-pulse stimulation.

**Results:** Individuals with CLBP had significantly higher rMT (decreased corticospinal excitability) and reduced ICF compared to controls. No significant differences were found in MEP amplitude and SICI.

**Conclusion:** These findings add to the growing body of evidence that CLBP is associated with deficits in intracortical modulation involving glutamatergic mechanisms.

**Significance:** This article reports chronic lower back pain is associated with changes in intracortical excitability, specifically reduced intracortical facilitation. Furthermore, the imbalance between facilitation and inhibition may be related to pain intensity. These findings may help clinicians in the treatment of chronic pain with an increased focus on using neuromodulation techniques, targeting maladaptive intracortical facilitation, as a potential therapeutic tool in chronic pain.

## Introduction

Chronic pain is associated with altered excitability in the motor cortex (Moseley & Flor, 2012; Parker et al., 2016). Motor control dysfunction is common across many chronic pain conditions, including Chronic Lower Back Pain (CLBP). There is increasing support that these deficits are associated with pain-related plasticity (altered intracortical and corticospinal excitability; Massé-Alarie & Schneider, 2016; Parker et al., 2016). Those with CLBP and fibromyalgia demonstrate overall reductions in corticospinal excitability (Strutton et al., 2005; Mhalla et al., 2010). However, the dearth of research in this field means that the specific mechanisms underlying altered excitability in CLBP remain unclear (Schabrun & Hodges, 2012).

Evidence for altered motor cortex excitability in CLBP is inconclusive (Chang et al., 2018). Strutton et al. (2005) reported that people with CLBP demonstrated significantly higher motor thresholds (decreased cortical excitability) compared to controls. Conversely, Massé-Alarie et al. (2012; 2017) reported no significant difference in motor threshold and MEP amplitude between CLBP and controls. Tsao et al. (2008) also reported no significant difference in motor threshold between people with recurrent LBP and controls. The lack of consistent findings in global excitability in CLBP has led to investigating if specific intracortical mechanisms may underlie changes in excitability in CLBP.

Short intracortical inhibition (SICI) and facilitation (ICF) are the main mechanisms underlying cortical plasticity (Lotze & Moseley, 2007). Studies in induced pain have reported changes in SICI and ICF during pain and after the removal of pain, suggesting that modulation of intracortical excitability is affected by pain perception (Brighina et al., 2011; Fierro et al., 2010). The restoration of normal SICI and ICF after the removal of pain suggests that the inability to restore normal intracortical excitability is associated with increased pain. This suggests SICI and ICF may be the key mechanisms associated with the maintenance of pain (Schabrun & Hodges, 2012).

Studies examining excitability in CLBP are limited. One study reported a decrease in corticospinal excitability in CLBP but did not examine intracortical mechanisms, SICI and ICF (Strutton et al., 2005). Massé-Alarie et al. (2016) reported reduced SICI in people with CLBP compared to controls. There were, however, no significant differences between short ICF or cortical silent period (Massé-Alarie et al., 2016). Massé-Alarie et al. (2017) reported no difference in SICI and short ICF between people with CLBP and controls. In comparison, decreased SICI has been reported in those with chronic hand pain (Lefaucheur et al., 2006), and fibromyalgia, where both SICI and ICF levels are reduced compared to controls (Mhalla et al., 2010). Although the pattern of change in these studies differs, altered motor cortex excitability is the common element of chronic pain.

A greater understanding of intracortical excitability in CLBP is required. It is unclear if people with CLBP have altered excitability and, if so, if this contributes to the persistence of CLBP. The present study sought to explore how intracortical mechanisms within the motor cortex may differ between people with CLBP and controls. The relationships between motor cortex excitability and pain-related measures were also examined.

## Methods

### Participants

Participants were recruited via convenience sampling to participate in a 6-week intervention study. The participants’ assessments at baseline form the data for this study. This study was approved by Curtin University ethics committee and all research was conducted in accordance with the Declaration of Helsinki. All participants provided written, informed consent. Inclusion in the study required a formal diagnosis of CLBP by a qualified health professional (General Practitioner or Physiotherapist) of at least 6 months (see Table 1 for demographic information and pain related information). Individuals were screened against Transcranial Magnetic Stimulation (TMS) inclusion criteria and screened for cognitive status using the Telephone Interview for Cognitive Status – 30 (TICS-30; score ≥ 18 for inclusion). Thirty-one participants met the initial inclusion criteria for participation. Eleven participants did not produce reliable MEPs and were removed from subsequent TMS analysis. Twenty participants were included in the final data set. Control participants were recruited based on age and gender-match to the CLBP participants (n = 18).

**Table 1.** Demographics, Pain Classification, and Treatment Engagement. *Note*. CLBP Classification = classification of chronic lower back pain based on Koes et al. (2001), Non-Specific = no radiographical injury at time of participation, Specific = Radiographical evidence, Pain Average = Average pain intensity one week prior to participation. Other = Acupuncture, Chiropractor, Massage). ^a^ = Percentage based on individuals taking medication.

|  | Total |  | CLBP |  | Control |  |
| --- | --- | --- | --- | --- | --- | --- |
|  | CLBP<br>(n = 20) | Control<br>(n = 18) | Males<br>(n = 11) | Females<br>(n = 9) | Males<br>(n = 12) | Females<br>(n = 6) |
| Age | 54.45 (15.89) | 53.83<br>(16.72) | 59.73<br>(15.87) | 48.00<br>(14.12) | 58.17<br>(16.47) | 45.17<br>(14.78) |
| Years of Education | 12.47 (3.23) | 14.06 (4.95) | 12.05 (2.66) | 12.99 (3.92) | 11.95 (2.67) | 13.58 (3.62) |
| Duration of Diagnosis (years) | 13.94 (13.12) | - | 18.98<br>(15.12) | 8.33 (7.92) | - | - |
| VAS Pain Average | 5.02 (2.07) | - | 4.64 (2.27) | 5.44 (1.87) | - | - |
| CLBP Classification |  |  |  |  |  |  |
| Non-Specific | 85% | - | 82% | 89% | - | - |
| Specific | 15% | - | 18% | 11% | - | - |
| Percentage taking Pain Medication | 75% | - | 55% | 100% | - | - |
| Anti-Inflammatory (Celebrex, Mobic,<br>Ibuprofen)* | 53% (n = 8) | - | 67% (n = 4) | 44% (n = 4) | - | - |
| Pain Killer (Over the counter and<br>prescription<sup>1</sup>)* | 67% (n = 10) | - | 50% (n = 3) | 78% (n = 7) | - | - |
| Benzodiazepine (Valium, Norflex)* | 27% (n = 4) | - | 17% (n = 1) | 33% (n = 3) | - | - |
| Anti-Depressants (Endep)* | 13% (n = 2) | - | 17% (n = 1) | 11% (n = 1) | - | - |
| Engaging in Physiotherapy | 50% | - | 36% | 67% | - | - |
| Past Surgery | 25% | - | 27% | 22% | - | - |
| Other Pain Management (Chiropractor) | 75% | - | 73% | 78% | - | - |
| Depression and Anxiety Disorder | 20% (n = 3) | 17% (n = 3) | 9% (n = 1) | 22% (n = 2) | 17% (n = 2) | 17% (n = 1) |
| Anti-Anxiety Medication* | 33% (n = 1) | 67% (n = 2) | 0% | 50% (n = 1) | 100% (n = 2) | 0% |

Acupuncture, Chiropractor, Massage). \* = Percentage based on individuals taking medication.<sup>1</sup> = 5 participants were taking prescription medication; Tramadol, Tapentadol, Lyrica and/or Codeine.

### Measures

Demographic and pain-related information were collected via self-report questionnaire. All CLBP participants completed TMS and clinical measures. Control participants only completed the TMS measures.

### Motor Cortex Excitability Measures

EMG signals were recorded from the left First Dorsal Interosseous (FDI) muscle using Ag – AgCl surface electrodes placed over the belly and tendon of the FDI. The stimulation procedures were conducted using TMS. TMS was applied using a figure-of-eight coil (90mm in diameter) that was connected to two Magstim 200 magnetic stimulators through a Bistim module (Magstim Company Limited, UK). The motor area corresponding to the left FDI muscle was located using the 10/20 International system for electrode placement (Trans Cranial Technologies, 2012). The coil was positioned over the optimal location to produce a MEP in the contralateral FDI at a 45-degree angle from the inter-hemispheric line, with the handle pointing towards the right-hand side, to stimulate current flow in a posterior to anterior direction.

To determine rMT, stimulation intensity started at 30% and was adjusted in 1% increments until the rMT was established. rMT was established as the lowest stimulus level that elicits MEPs of greater than 50μV in 3 of 5 trials with the average of the five MEPs also greater than 50μV. To determine the recruitment curve stimulation intensity, the stimulation intensity was adjusted in 1% increments, until a mean MEP of 1mV was produced in eight trials. The recruitment curve consisted of stimulation at 90%, 100% (1mV), 110%, 120%, and 130% of the intensity required to produce the 1mV MEP. The order of administration was randomised. Eight pulses were delivered for each intensity level and were averaged to attain a mean MEP amplitude. The mean MEP amplitude for each intensity level was normalised against the participant’s 1mV response.

SICI and ICF were measured using the paired-pulse protocol developed by Kujirai et al. (1993) SICI and ICF were assessed using a subthreshold conditioning pulse set to 80% of rMT, followed by a suprathreshold test pulse set at 120% of rMT. The interstimulus interval was set to 3ms for SICI and 10ms for ICF. Fifteen trials were recorded at each interstimulus interval, and fifteen single unconditioned test pulses were also recorded (set at 120% rMT), with an 8 second interval between each trial. The order of administration was randomised. The fifteen trials for each interstimulus interval were averaged to attain a mean MEP amplitude. The mean MEP amplitude for each interstimulus interval was normalised against the participant’s mean unconditioned pulse.

### Clinical Measures

#### Pain Intensity

The Short-Form McGill Pain Questionnaire (SF-MPQ) contains a 10 cm Visual Analogue Scale (VAS) and was used to assess average pain intensity in CLBP (Melzack, 1987). Participants were required to mark the line at the spot they feel applied to their level of pain (Hawker et al., 2011).

#### Disability

The Roland-Morris Disability Questionnaire (RMDQ) assessed the level of disability in CLBP (Roland & Morris, 1983). The RMDQ consists of 24 items assessing the impact of CLBP across multiple domains (mobility, daily activities, sleeping, mood, and appetite).

#### Pain Catastrophising

The Pain Catastrophizing Scale (PCS) assessed the presence of pain catastrophising in individuals with CLBP (Sullivan et al., 1995). The PCS consists of 13 items assessing rumination, magnification, and helplessness.

### Statistical Analysis

All analyses were conducted using R software (v3.5.1; R Foundation for Statistical Computing, Vienna, Austria). All trials were visually inspected and peak to peak MEP amplitudes were manually marked. Trials were excluded from analysis if visual inspection indicated noise, artifacts, or voluntary contraction, which obscured the detection of MEP amplitude, was present in the EMG signal. Robust ANOVAs and independent samples T-tests were conducted using WRS2 package (Mair & Wilcox, 2020). Recruitment curve data was analysed using robust two-way ANOVA, pbad2way function. Group differences in rMT, SICI, ICF, and were analysed using Yuen-Welch robust t-test with bootstrapping, yuenbt function (nboot = 10,000). Correlations were analysed using robust correlation, pbcor function.

## Results

There was a significant difference in rMT between the CLBP and the control group (T_pb_ = −3.00 [−12.16, −2.75], p = 0.004, trimmed mean difference = −7.56; see Figure 1). The robust ANOVA revealed a significant main effect of Intensity (p < .001; MEP ratios increased with Intensity) but there was no significant main effect of Group (p = .132) or two-way interaction (p = .816), indicating that the pattern of corticospinal excitability at-rest did not differ between CLBP and the control group (See Figure 2).

**Figure 1.**
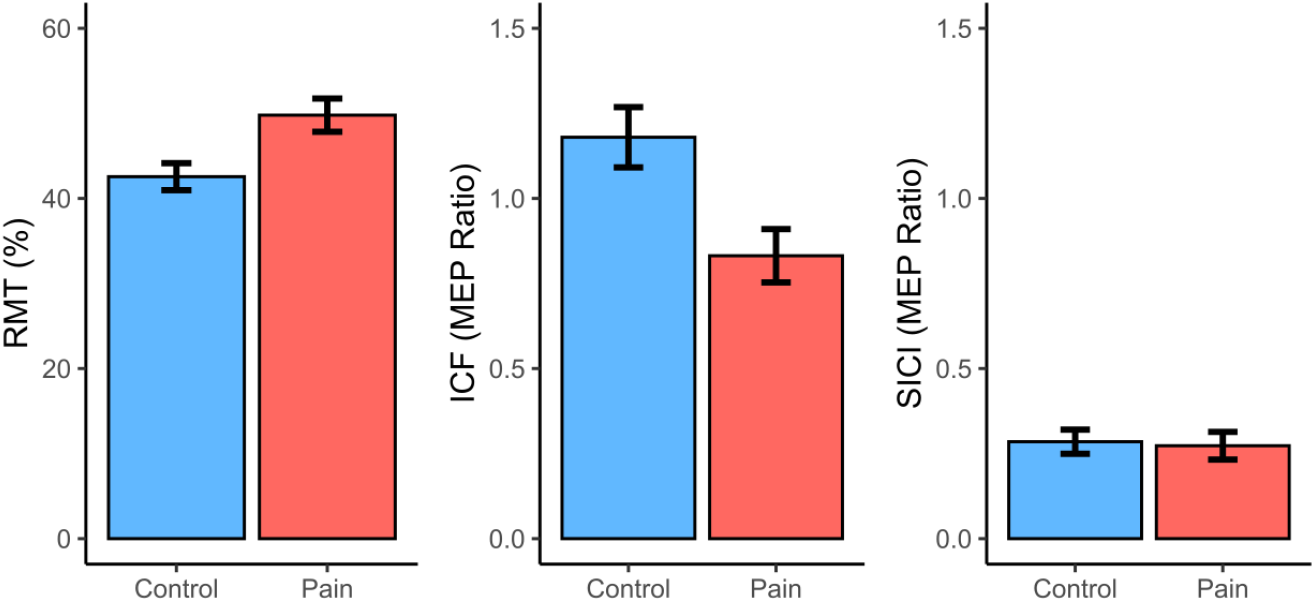
Average rMT, ICF, and SICI MEP Ratio for CLBP and Control Group.

**Figure 2.**
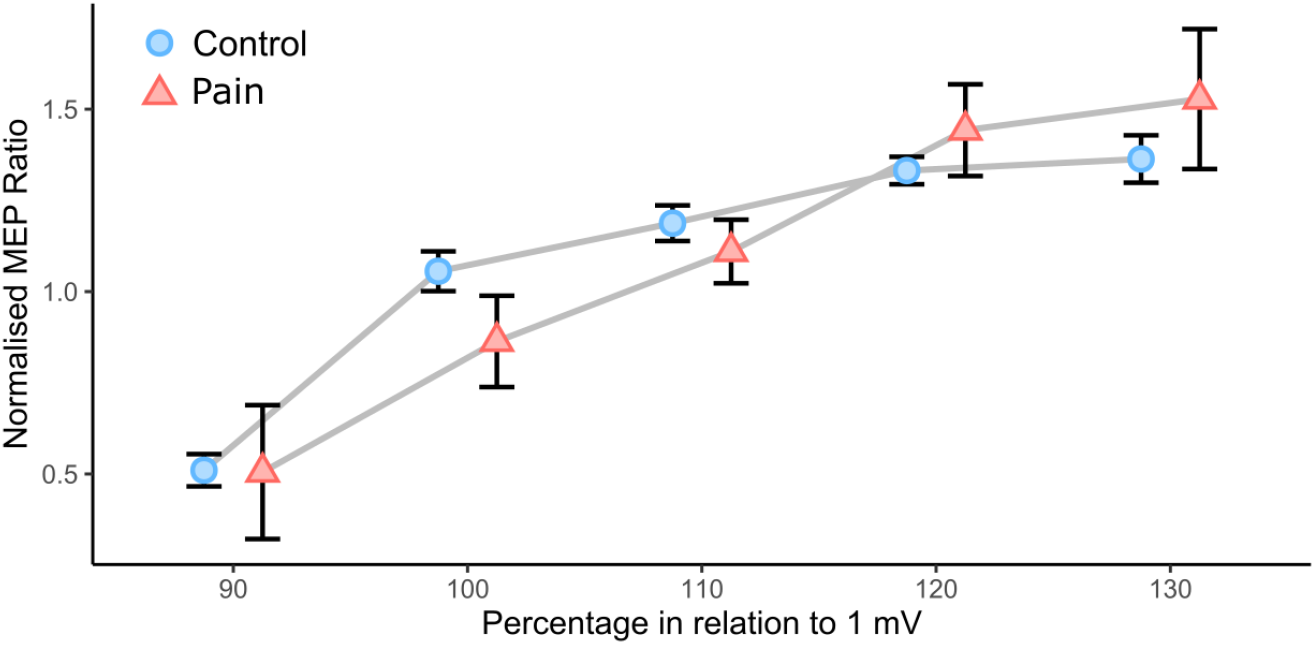
Recruitment Curve with Normalised MEP Ratio at 90 – 130% Stimulation for CLBP and Control Group. 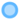 Control 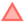 Pain

There was no significant difference in SICI between groups (T_pb_ = 0.47 [−0.10, 0.16], p = 0.632, trimmed mean difference = 0.03; see Figure 1). ICF was significantly reduced in the pain group (T_pb_ = 2.61 [0.09, 0.61], p = 0.016*, trimmed mean difference = 0.35). There was no significant correlation between pain intensity and ICF (r_pb_ = 0.21, T_pb_ = 0.90, p = .379) or RMT (r_pb_ = 0.03, T_pb_ = 0.14, p = .891). Correlations between ICF and other pain measures (PCS total and RMDQ total) were non-significant.

### Post Hoc Analysis

As cortical excitability reflects the balance between excitation and inhibition, post hoc analysis was conducted to determine if the balance between SICI and ICF was related to pain. To determine the difference score between SICI and ICF, SICI MEP amplitudes were subtracted from ICF MEP amplitudes ([MEPSICI/MEPcontrol] - [MEPICF/MEPcontrol). The correlation between pain intensity and SICI-ICF difference approached significance (r_pb_ = 0.44, T_pb_ = 2.10, p = .050*; See Figure 3). Inspection of the posterior distribution (See Figure 4) Bayesian General Linear Models showed a positive estimate median of 0.06 (95% CI: [0.00, 0.12]) for each unit increase in the McGill VAS, with positive directional probability of 97.02%, 86% probability of significance, and a 22.27% of being large (> 0.08). Correlations between SICI-ICF difference and other pain measures (PCS total, and Roland Morris Total) were all non-significant.

**Figure 3.**
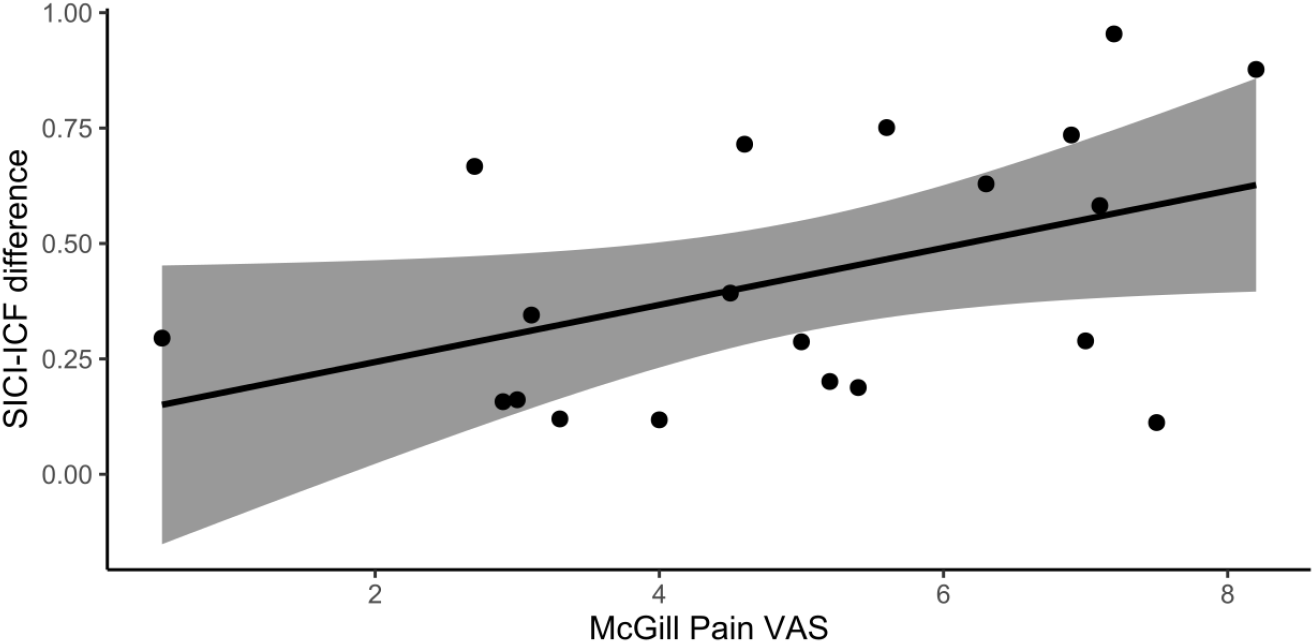
Correlation between pain intensity (McGill VAS) and SICI-ICF difference (positive = inhibition).

**Figure 4.**
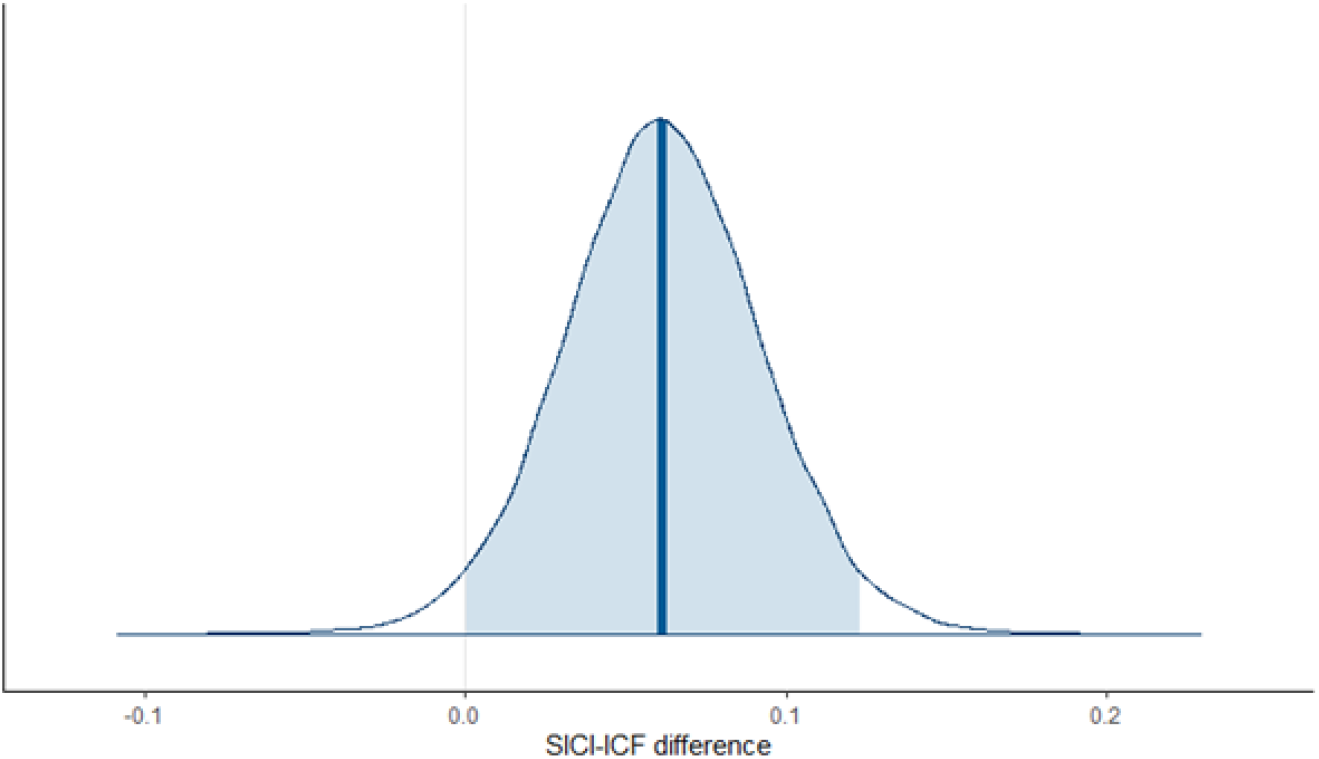
Bayesian posterior distribution of SICI-ICF difference for each unit increase in pain intensity (McGill VAS; 95% CI).

### Discussion

The present study explored motor cortex excitability in people with CLBP, compared to pain-free, age, and gender-matched controls. Previous research has focused on varying forms of chronic pain (including CLBP), producing conflicting results. The present study revealed that CLBP is associated with changes in motor cortical excitability. The CLBP group demonstrated higher rMT and reduced ICF compared to controls, but there were no differences in SICI between the two groups. Differences in rMT and ICF were not associated with pain intensity, duration, pain sensation, use of medication, or disability in those with CLBP.

The evidence for change in motor cortex excitability in CLBP is inconclusive (Chang et al., 2018). Strutton et al. (2005) reported that individuals with CLBP had a significantly higher motor threshold, which is typically indicative of decreased global excitability (Mhalla et al., 2010; Schoenen et al., 2008). Conversely, Massé-Alarie et al. (2012) reported no significant difference in motor threshold between people with CLBP and controls. The present finding of higher rMT in CLBP is consistent with Strutton et al. (2005) and suggests that the cortical system is less excitable compared to controls. rMT is a marker for the trans-synaptic excitability of corticospinal output neurons (Perez & Cohen, 2009), with a higher threshold indicative of reduced trans-synaptic excitability. Hypoexcitability of the corticospinal tract is thought to indicate reduced activity of motoneurons, and as such, less likely to respond to motor commands. This results in dysfunction in voluntary motor control (Powers et al., 2014). In accord with this, the present finding suggests that the balance of neurochemicals in the motor cortex may be compromised in those with CLBP.

The present study found that ICF was significantly decreased in CLBP compared to controls, a finding consistent with fibromyalgia research (Mhalla et al., 2010; Salerno et al., 2000). ICF involves excitatory glutamatergic interneurons, and reduced ICF is indicative of hypoexcitability of motor circuity (Powers et al., 2014). This is the first study to show decreased ICF in people with CLBP compared to pain-free controls. These findings are consistent with a motor cortex mapping study that reported reduced corticospinal excitability is associated with smaller map volume in the motor cortex in severe CLBP (Schabrun et al., 2017). Neurochemical N-Acetylaspartate, which acts on excitatory glutamate receptors, have also been shown to be reduced in CLBP compared to pain-free controls (Sharma et al., 2012). Sharma et al. (2012) suggested that a reduction in N-Acetylaspartate may underlie functional motor cortex changes in CLBP. This provides further support for the hypoexcitability of the motor cortex in CLBP.

Hypoexcitability has been reported for other forms of chronic pain, including arthritis (Salerno et al., 2000) and fibromyalgia (Mhalla et al., 2010; Salerno et al., 2000). Although hypoexcitability may result from both spinal and supraspinal mechanisms, Mhalla et al. (2010) and Schoenen et al. (2008) suggested that hypoexcitability (as indicated by higher motor threshold) involves supraspinal mechanisms, as opposed to spinal mechanisms (Mhalla et al., 2010; Schoenen et al., 2008). The involvement of supraspinal mechanisms was supported by a lack of change in the H-reflex and dysfunctional motor control in fibromyalgia (Mhalla et al., 2010; Schoenen et al., 2008). Given the similar findings in the present study and frequently reported motor dysfunction in previous CLBP studies, it seems reasonable to suggest that hypoexcitability in people with CLBP may also involve supraspinal mechanisms. Establishing the involvement of supraspinal mechanisms in hypoexcitability in CLBP may be of clinical importance as there is evidence that cortical disruption contributes to, and/or maintains, chronic pain (Meier et al., 2019; Moseley & Flor, 2012).

The present findings lend support to the use of neuromodulation techniques, such as non-invasive brain stimulation, as a potential therapeutic tool in the treatment of CLBP. For instance, transcranial Direct Current Stimulation (tDCS) modulates the spontaneous firing of neurons and is able to reverse maladaptive changes in plasticity (Malavera et al., 2015). Research has investigated the use of tDCS in combination with pharmacological and physical rehabilitation in CLBP. Pinto et al. (2018) reported that anodal tDCS combined with behavioural therapy or nerve stimulation in individuals with chronic pain led to the greatest reduction in pain, compared to individual therapies or controls. In CLBP, the combination of tDCS and nerve stimulation increased motor cortex excitability, normalised motor cortex organisation, and reduced pain (Hazime et al., 2017; Schabrun et al., 2018).

In comparison to other CLBP studies, the present findings did not indicate any changes in inhibition (SICI). Strutton et al. (2005) reported decreased GABA inhibition in people with CLBP, as measured by the cortical silent period, but did not investigate any changes in SICI or ICF. Conversely, Massé-Alarie et al. (2016) reported no difference in GABAB inhibition in people with CLBP but did report a decrease in GABA_A_ SICI. These inconsistent results may relate to the TMS paradigm used to measure inhibition. To the best of our knowledge, this is the first study to measure SICI and ICF from the left FDI in CLBP. The FDI is a widely used, easy to target muscle for measuring motor cortex excitability and provides consistent results (Groppa et al., 2012). Furthermore, altered concentration of GABA and glutamate may reflect more of a global alteration in cortical excitability that is not restricted to the region representing the FDI (Parker et al., 2016).

Given the involvement of motor cortical structures in movement planning and execution (Schabrun et al., 2017), it is likely that changes in motor cortex excitability are associated with decreased motor control in people with CLBP. Several studies have reported motor control dysfunction in CLBP and chronic pain, including increased disability (Strutton et al., 2005), fatigue (Schabrun et al., 2017), depression, and catastrophising (Mhalla et al., 2010). Motor control dysfunction and reduced intracortical excitability have also been reported in Fibromyalgia. Mhalla et al. (2010) reported that the intensity of fatigue in fibromyalgia was correlated with decreased ICF, while depression and catastrophising was associated with decreased SICI. Furthermore, chronic pain studies have reported the restoration of normal SICI and ICF levels when pain was removed (Antal et al., 2010; Fregni et al., 2006a; Fregni et al., 2006b). While the present study did not reveal significant relationships between pain intensity, disability, catastrophising, and specific measures of cortical excitability, post hoc analysis revealed a relationship between the balance of excitability and inhibition, and pain intensity. Results showed that an increase in inhibition was associated with increased pain intensity. These results suggest that the relationship between cortical excitability and pain cannot be simply explained by a gross or singular measure (such as ICF) in isolation but may be the result of the interaction between multiple mechanisms. While the finding that increased pain is associated with SICI appears incongruent with the overall finding of decreased ICF, it is possible that reduced excitability may be regulated by an increase in inhibition to keep the nervous system within a functional, dynamic range (Filmer et al., 2019). As a result, pain might be a by-product of neural dysfunction when excitation/inhibition mechanisms are not balanced.

There are a few limitations that must be acknowledged. The use of analgesic medications can influence cortical excitability. Benzodiazepines are reported to significantly increase SICI, as benzodiazepines increase GABA_A_ inhibitory transmission (Di Lazzaro et al., 2006). In contrast, ICF is decreased by GABA_A_ receptor agonists (such as benzodiazepines), and N-methyl-D-aspartate receptor agonists (synthetic opioids; Schwenkreis et al., 1999). While medication use was recorded in the present study, frequency of use and dosage was not documented. Although a limitation, correlations revealed that there was no relationship between medication use and any motor cortex excitability measures. Given this, and as analgesics were not associated with changes in cortical excitability and modulation in fibromyalgia (Mhalla et al., 2010), medication use is not thought to underlie the findings of the present study.

These findings add to the growing body of evidence that CLBP is associated with changes in intracortical excitability. While it appears changes in ICF may contribute to the pathophysiology of CLBP, future studies should examine if these changes are directly responsible for CLBP symptoms, or an indirect result of the interaction between multiple mechanisms. Nonetheless, these results may have direct clinical applications in terms of the use of neuromodulation techniques for the treatment of CLBP. Previous research has suggested that functional improvements in chronic pain are related to the restoration of altered intracortical excitability (Lefaucheur et al., 2006; Mhalla et al., 2010). Given the present findings show that altered intracortical excitability occurs in CLBP, the use of neuromodulation techniques may be of clinical significance for rehabilitation and treatment of people with CLBP.

## Authorship

**EJC: S**ubstantial contributions to conception and design; acquisition of data, analysis and interpretation of data; drafting the article, and final approval of the version to be published.

**WM:** Substantial contributions to analysis and interpretation of data; revising article critically for important intellectual content, and final approval of the version to be published.

**ATN:** Substantial contributions to analysis and interpretation of data; revising article critically for important intellectual content, and final approval of the version to be published.

**NG:** Substantial contributions to conception and design; revising article critically for important intellectual content, and final approval of the version to be published.

**AML: S**ubstantial contributions to conception and design; acquisition of data, analysis and interpretation of data; revising article critically for important intellectual content, and final approval of the version to be published.

## Notes

<u>Declarations of interest:</u> None of the authors have potential conflicts of interest to be disclosed.

### Competing Interest Statement

The authors have declared no competing interest.

### Summary of Updates

Revised Table 1 for additional clarity on number of participants taking medication.

